# Effects of the estrous cycle and ovarian hormones on cue-triggered motivation and intrinsic excitability of medium spiny neurons in the Nucleus Accumbens of female rats

**DOI:** 10.1101/669804

**Authors:** Yanaira Alonso-Caraballo, Carrie R. Ferrario

## Abstract

Naturally occurring alterations in estradiol influence food intake in females. However, how motivational responses to food cues are affected by the estrous cycle or ovarian hormones is unknown. In addition, while individual susceptibility to obesity is accompanied by enhanced incentive motivational responses to food cues and increased NAc intrinsic excitability in males, studies in females are absent. Here, we examined basal differences in intrinsic NAc excitability of obesity-prone vs. obesity-resistant females and determined how conditioned approach (a measure of cue-triggered motivation), food intake, and motivation for food vary with the cycle in naturally cycling female obesity-prone, obesity-resistant, and outbred Sprague-Dawley rats. Finally, we used ovariectomy followed by hormone treatment to determine the role of ovarian hormones in cue-triggered motivation in selectively-bred and outbred female rats. We found that intrinsic excitability of NAc MSNs and conditioned approach are enhanced in female obesity-prone vs. obesity-resistant rats. These effects were driven by greater MSN excitability and conditioned approach behavior during metestrus/diestrus vs. proestrus/estrus in obesity-prone but not obesity-resistant rats, despite similar regulation of food intake and food motivation by the cycle in these groups. Furthermore, estradiol and progesterone treatment reduced conditioned approach behavior in obesity-prone and outbred Sprague-Dawley females. To our knowledge, these data are the first to demonstrate cycle- and hormone-dependent effects on the motivational response to a food cue, and the only studies to date to determine how individual susceptibility to obesity influences NAc excitability, cue-triggered food-seeking, and differences in the regulation of these neurobehavioral responses by the cycle.

## I. INTRODUCTION

Naturally occurring fluctuations in ovarian hormones influence food consumption and food preference in both humans and rodents (Eckel, 2011; Hirschberg, 2012). For example, in rats, food intake is lower during estrus compared to metestrus and diestrus (Tarttelin Gorski, 1971). This effect of the cycle on food intake is mediated in part by central actions of estrogens (see Asarian and Geary, 2013 for review). Consistent with this, the removal of ovarian hormones via ovariectomy produces significant increases in food intake that are accompanied by weight gain (Blaustein Wade, 1976; Yu et al., 2011). This increase can be normalized by treating the ovariectomized rats with estradiol (Tarttelin Gorski, 1971; Wade, 1972; Asarian et al., 2002, 2006; Yu et al., 2011). Thus, ovarian hormones influence food intake.

In overweight women, stimuli that predict food availability (i.e., food cues) elicit stronger activations in brain regions that influence motivation including the Nucleus Accumbens (NAc; Stoeckel et al., 2008) compared to lean women. Furthermore, the magnitude of food cue elicited NAc activations predicts future weight gain in healthy weight females and future inability to lose weight after obesity (Murdaugh et. al 2012, Demos et al. 2012). These human studies suggest that enhanced neurobehavioral responses to food cues may promote the development of obesity. However, identifying pre-existing differences is not feasible in human studies and there is limited understanding of how ovarian hormones influence brain mechanisms governing cue-triggered food seeking (see Alonso-Caraballo et al., 2018 for additional discussion).

Consistent with a role for pre-existing differences, pre-clinical studies from our lab have shown that motivational responses to food cues are stronger in obesity-prone compared to obesity-resistant male rats (Derman and Ferrario, 2018; Robinson et al., 2015). These behavioral differences in males are mediated by the NAc (Derman and Ferrario, 2018) and intrinsic excitability of medium spiny neurons (MSNs) in the NAc is enhanced in obesity-prone vs. obesity-resistant males (Oginsky, et al., 2016). The risk for obesity is greater in females compared to males, and ovarian hormones influence NAc activity and plasticity (Cyr et al., 2001; Le Saux et al., 2006; Peterson et al., 2015; Cao et al., 2018). However, it is not known whether similar enhancements in motivational responses to food cues are also found in female obesity-prone vs. obesity-resistant rats. Furthermore, neural activations in response to monetary rewards or to visual food cues vary with the menstrual cycle in women, with higher activations observed in corticolimbic areas during the luteal phase compared to the follicular phase (Dreher et al., 2007; Frank et al., 2010; Arnoni-Bauer et al., 2017). This suggests that motivational responses to food cues may be influenced by ovarian hormones; however, no studies to date have directly examined this possibility.

In the current study, we first examined basal differences in intrinsic excitability of NAc MSNs of obesity-prone vs. obesity-resistant females across the cycle. Next, we determined how motivational responses to Pavlovian food cues and motivation for food itself vary with the cycle in naturally cycling female obesity-prone and obesity-resistant rats. Finally, we used ovariectomy followed by hormone treatment to determine the role of ovarian hormones in cue-triggered motivation in selectively bred and outbred female rats.

## II. MATERIALS AND METHODS

### Subjects

Female selectively-bred obesity-prone (OP) and obesity-resistant (OR) rat lines were originally developed by Barry Levin (Levin et al., 1997) and were bred in house. Female outbred Sprague-Dawley rats were purchased from Charles River Breeding Labs (Portage, MI). Rats were housed on a reverse 12-h light/dark cycle, had free access to food and water throughout, and were group housed unless otherwise noted. For studies involving ovariectomy, estrogen-free bedding (7090 Teklad sanichips) and estrogen-free chow (Harlan 2916: 3 kcal/g; 4% fat; 16.4% protein; 48.5% protein; % of caloric content) were used. Rats were weighed 3-4 times per week unless otherwise specified. Procedures were approved by The University of Michigan Committee on the Use and Care of Animals in accordance with AAALAC and AVMA guidelines. See also https://sites.google.com/a/umich.edu/ferrario-lab-public-protocols/ for additional details.

### Monitoring the Estrous Cycle

Estrous cycle phase was determined by daily observations of vaginal epithelial cell cytology, precopulatory, and copulatory behaviors (Marcondes et al., 2002). Epithelial cells were collected daily by vaginal lavage (during the dark phase) and visualized using an inverted light microscope (Olympus CKX53) under bright-field. Estrous cycle cell pictures were taken at 20x magnification with an EVOS XL Cell Imaging System (Thermo Fisher Scientific). Cell morphology was then used to determine cycle phase with metestrus characterized by a mix of lymphocytes, cornified cells and epithelial nucleated cells, diestrus by lymphocytes and a little to any epithelial nucleated cells, proestrus by nucleated cells that form sheets, and estrus by masses of large cornified cells that lack nuclei (see Fig. 2A). Body weight, food intake, precopulatory and copulatory behaviors (such as ear wiggling, darting, and lordosis) were also used to further verify the estrous cycle phase.

### Behavioral training and testing

For all behavioral studies, rats were trained and tested hungry (chow removed from cages, 5-6 hrs prior to training/testing and replaced following training/testing). All training and testing occurred in standard operant boxes (Med Associates, St. Albans City, VT) housed within sound attenuating chambers. Testing occurred at the same time each day, approximately 6-8 hours after the start of the dark cycle. Instrumental procedures: To evaluate motivation for food in obesity-prone and obesity-resistant females, we used instrumental training followed by progressive ratio testing. First, rats underwent two magazine training sessions in which 20 sucrose pellets (45 mg TestDiet; cat.) were delivered into the food cup on a variable interval of 60 seconds (VI 60). Next, they were trained to press one lever (active) that resulted in the delivery of one sucrose pellet. Pressing a second lever (inactive) had not consequences but was recorded. No discrete cues were paired with pellet delivery. Left/right position of the active and inactive levers relative to the food cup was counterbalanced. Rats were trained in three sessions in which each response on the active lever resulted in delivery of a food pellet (i.e., fixed ratio 1 [FR1], 60min/ session or until 50 pellets were earned). This was followed by three sessions in which five responses on the active lever were required to receive one food pellet (FR5, 60min/session or until 50 pellets were earned). Progressive ratio (PR) testing was then used to determine motivation to obtain food. During PR testing the number of lever presses required to obtain each subsequent sucrose pellet increased exponentially (5e(0.2*delivery+1)-5; adapted from (Richardson and Roberts, 1996). The PR session ended when rats did not meet the next ratio requirement within 60 minutes (i.e., breakpoint). The number of active and inactive lever presses, pellets earned, and final break point achieved were recorded.

### Pavlovian procedures

First rats underwent two magazine training sessions in which 20 sucrose pellets were delivered into the food cup on a VI 60 schedule. During Pavlovian conditioning sessions, one auditory cue (CS+; 2min) was paired with the delivery of 4 sucrose pellets (US), whereas a second auditory cue (CS-; 2min) was never paired with sucrose (CS+/CS-tone or white noise counterbalanced). Sucrose pellets were delivered on a VI of 20 seconds during CS+ presentation. A total of 4 CS+ and 4 CS-trials separated by a variable 5-min intertrial-interval (ITI; range 2-7 min) were given per session (5-8 sessions, 60 min per session, 1 session/day). Subsequent testing was identical to initial conditioning except that no sucrose was given. Food cup entries during the ITI, CS+ and CS-periods were recorded throughout.

### Electrophysiology

Whole-cell patch clamp recordings of MSNs in the NAc core were conducted in naturally cycling obesity-prone and obesity-resistant rats (P70-P80) during the metestrus/diestrus or proestrus/estrus phases using established procedures (e.g., Oginsky et al., 2016). Briefly, rats were anesthetized with chloral hydrate (400mg/kg, i.p.) prior to slice preparation, brains were rapidly removed and placed in icecold oxygenated (95% O2 5% CO2) aCSF containing (in mM): 125 NaCl, 25 NaHCO3, 12.5 glucose, 1.25 NaH2PO4, 3.5 KCl, 1 Lascorbic acid, 0.5 CaCl2, 3 MgCl2, pH 7.4, 305 mOsm. A vibratome (Leica Biosystems, Buffalo Grove, IL, USA) was used to make 300 m coronal slices containing the NAc. Before recording, slices were allowed to rest in oxygenated aCSF for at least 40 minutes at 35°*C*, followed by a 10-minute recovery time at room temperature. For the recording aCSF, CaCl2 was increased to 2.5 mM and MgCl2, was decreased to 1mM. Patch pipettes were pulled from 1.5 mm borosilicate glass capillaries (WPI, Sarasota, FL; 4-7 M Ω resistance) with a horizontal puller (model P97, Sutter Instruments, Novato, CA, USA) and filled with a solution containing (in mM): 130 K-gluconate, 10 KCl, 1 EGTA, 2 Mg2+-ATP, 0.6 Na+-GTP and 10 HEPES, pH 7.45, 285 mOsm. Recordings were conducted in the presence of the GABAA receptor antagonist, picrotoxin (50*µM*). MSNs in the NAc core were identified based on resting membrane potential and action potential firing in response to current injection (−200 to +200pA, 25pA increments, 500ms). For data analysis, only cells with a leak of less than 50pA and access resistance of less than 30pA were used. These cell parameters were recorded before at the start and end of data collection and only cells with less than 20% change between the start and end of recordings were included in analyses. I/V relationships were determined by calculating the difference between the baseline voltage and the voltage 200ms after initial current injections. Input resistance was determined by the change in voltage from −50pA to +50pA current injections. The number of action potentials elicited by each depolarizing current injection were used to determine neuronal excitability. Rheobase was defined as the minimum amount of current injection to elicit an action potential. The maximum second derivative method (Sekerli et al., 2004) was used to determine the action potential threshold.

### Ovariectomy and hormone treatment

Rats were bilaterally ovariectomized under isoflurane anesthesia (2.5-5%, inhalation) via a single back incision. Before surgery rats were given the analgesic Carprofen (3mg/kg, s.c.; Rimadyl, Henry Schein). The incision was closed using dissolvable sutures internally (Reli Redi Gut Chromic Suture), and 9 mm wound clips externally (Reflex 9 mm wound clips, VWR international). In addition, carprofen was given again 48 hours after surgery. Rats were allowed to recover for at least 10 days prior to any behavioral testing. Vaginal lavages were taken beginning 7 days after surgery and throughout the remainder of the study.

The effect of three different hormone treatments were evaluated. Hormone treatment consisted of subcutaneous injections of 17*β*-estradiol benzoate (estradiol; Sigma-Aldrich, E8515) for two consecutive days (5*µg* in 0.1ml of peanut oil, 24 hrs apart), followed by a single progesterone injection on the third day (500*µg* in 0.1ml of peanut oil, Sigma-Aldrich, P0130). Doses and treatment regimen were based on (Cummings Becker, 2012 and Yu et. al., 2011). For repeated hormone treatment this cycle was repeated 3 or 4 times (see additional details below). Controls were given the same number of vehicle injections (0.1ml of peanut oil, s.c.). For studies of estradiol or progesterone alone, the same total number of injections were given, with progesterone or estradiol injection replaced by vehicle as appropriate. Finally, for all studies involving progesterone, the final progesterone injection was given 4-6 hrs prior to testing (see also timelines in Figs. 6A and 7A, D).

### Statistics and data analysis

Two-tailed t-tests, two-way or three-way ANOVAs and Sidak’s post-hoc multiple comparisons were used (Prism 8, GraphPad, San Diego, CA). Electrophysiology data were analyzed using both Clampfit 10.4 (Molecular Devices) and MATLAB (R2018b, Mathworks Software).

## A. Experiments

### Experiment 1: Verification of female obesity-prone and obesity-resistant phenotypes and effects of the cycle on food intake and motivation for food

We first verified differences in body weight, adiposity and food intake between obesity-prone (OP) and obesity-resistant (OR) females and evaluated effects of the cycle on food intake and motivation for food (Levin Govek, 1998). Rats were maintained on either standard lab chow (Lab Diet 5001: 4kcal/g; 4.5% fat, 23% protein, 48.7% carbohydrates; % of caloric content) or a junk-food diet (JF) made in house (Robinson et al., 2015; Oginsky et al., 2016). The junk-food diet was a mash of Ruffle potato chips (40g), Chips Ahoy (130g), Nesquik (130g), Jiff peanut butter (130g), Lab Diet 5001 (200g) and 180ml of water (19.6% fat, 14% protein, and 58% carbohydrates; 4.5 kcal/g). Before junk-food diet exposure, body composition was determined by nuclear magnetic resonance spectroscopy (NMR; Minispec LF90II, Bruker Optics) conducted by the University of Michigan Animal Phenotyping Core in female obesity-prone (n = 16) and obesity-resistant (n = 16) rats. The rats were then assigned to chow or junk-food groups (counter balanced by initial weight and fat mass within obesity-prone and obesity-resistant groups) in order to verify weight-gain phenotypes (n = 8 per group: OP-chow, OP-JF, OR-chow, and OR-JF). Food intake and body weight were monitored throughout the 4 weeks of junk-food vs. chow consumption and NMR measures were made again at the end of this period.

In a separate set of rats, we determined how food intake varied across the estrous cycle in naturally-cycling obesity-prone and obesity-resistant rats. Home cage food intake, body weight, and estrous cycle phase were measured for 13 days in singly-housed naturally cycling obesity-prone (n = 6) and obesity-resistant (n = 6) rats and estrous cycle phase was determined as described above. Finally, we also determined estrous cycle effects on motivation to work for food using progressive ratio testing. Naturally-cycling obesity-prone (n = 10) and obesity-resistant (n = 10) rats were trained to lever press for a sucrose pellet and motivation to obtain this food was then determined using the progressive ratio procedure described above. In order to determine potential differences in break point across the cycle, all rats were given 8 PR test sessions and data were analyzed according to cycle phase during testing. This allowed us to make within subject comparisons of motivation to work for food during metestrus/diestrus vs. proestrus/estrus phases.

### Experiment 2: Differences in NAc core MSN intrinsic excitability between obesity-prone and obesity-resistant female rats

Potential differences in intrinsic excitability of MSNs in the NAc core between obesity-prone and obesity-resistant rats were determined in naturally cycling females. Estrous cycle phase was monitored for 5-7 days as described above and reconfirmed 1 hour before slice preparation and electrophysiological recordings (OP-M+D n = 6 rats, 22 cells; OP-P+E n = 5 rats, 13 cells; OR-M+D n = 5 rats, 13 cells; OR-P+E n = 5 rats, 15 cells).

### Experiment 3: Modulation of cue-triggered motivation by the estrous cycle in obesity-prone and obesity-resistant rats

We determined whether there were differences in cue-triggered motivation (as measured by conditioned approach) between obesity-prone and obesity-resistant rats and the role of the estrous cycle in modulating this behavior. Naturally-cycling obesity-prone (n = 31) and obesity-resistant (n = 31) rats underwent 5 sessions of Pavlovian training followed by a single session of testing in extinction conditions. Vaginal lavages were taken after the end of each session to confirm cycle phase on the day of testing.

### Experiment 4: Effects of single and repeated estradiol-progesterone treatment on cue-triggered motivation in obesity-prone female rats

Here we determined the effect of estradiol and progesterone treatment on cue-triggered motivation in obesity-prone rats (obesity-resistant rats were not included because there were no effects of the cycle on cue-triggered motivation in this group, see results below). Naturally cycling obesity-prone rats (n = 20) underwent 8 sessions of Pavlovian training followed by ovariectomy and 10 days of recovery as described above. Some rats were then given one cycle of estradiol and progesterone treatment (see above) followed by a single test in extinction conditions (n = 10), while others received vehicle treatment prior to the extinction test. Next, rats in the vehicle group were treated with 3 cycles of hormone treatment while the rats that initially received one cycle of treatment were given repeated vehicle injections before undergoing an additional test in extinction conditions.

### Experiment 5: Effects of repeated hormone treatment on cue-triggered motivation in outbred female rats

Here we determined whether repeated hormone treatment with both estradiol and progesterone, or repeated treatment with each hormone alone were sufficient to alter cue-triggered motivation in outbred Sprague Dawley female rats. To examine effects of dual hormone treatment, naturally cycling outbred rats (n = 20) were trained and ovariectomized as described above. After recovery from surgery, they were assigned to vehicle (n = 10) or repeated hormone treatment (n = 10) groups, counter-balanced by post-operative weight. They received 4 consecutive cycles of either vehicle or hormone treatment followed by a single extinction test (see Fig. 7A). To assess the effects of each hormone on its own, a separate set of naturally-cycling outbred rats (n = 40) underwent 8 sessions of Pavlovian training followed by ovariectomy and recovery as described above; 2 rats were excluded from studies due to complications during ovariectomy surgery. Rats were then assigned to vehicle (Veh; n = 9), estradiol alone (E; n = 10), progesterone alone (P; n = 10) or estradiol and progesterone treated groups (E+P; n = 9), counterbalanced by post-operative weight. Body weight and food intake were measured throughout the hormone injection cycles.

## III. RESULTS

### Experiment 1a: Verification of female obesity-prone and obesity-resistant phenotypes

Figure 1 shows weight, adiposity and food intake of obesity-prone and obesity-resistant female rats at baseline (Fig. 1A-C) and after 4 weeks of junk-food exposure (Fig. 1D-F). Although body weight was similar between groups (Fig. 1A), fat mass was significantly greater in female obesity-prone vs. obesity-resistant groups at baseline (Fig. 1B: *t*_30_=3.593, p < 0.01). This was accompanied by significantly lower lean mass in obesity-prone vs. obesity-resistant groups (Fig. 1C: *t*_30_ = 3.642, p < 0.01). After 4 weeks of free access to junk-food or chow, obesity-prone rats remained heavier than obesity-resistant rats (Fig. 1D: Two-way ANOVA main effect of group: *F*_(1,28)_=9.05, p < 0.01, no diet x group interaction). Measurements of adiposity revealed that junk-food consumption significantly increased fat mass in both groups (Fig. 1E: Two-way ANOVA main effect of diet: *F*_(1,28)_=18.09, p < 0.001) and reduced lean mass (Fig. 1F: Two-way ANOVA main effect of diet: *F*_(1,28)_=9.78, p<0.01) compared to chow-fed groups. However, as expected, the effect of junk-food on fat mass was more pronounced in obesity-prone groups (Fig. 1E: Average change from chow OP: 16.3 ± 4.2; OR: 9.4 ± 1.8).

**FIG. 1:**
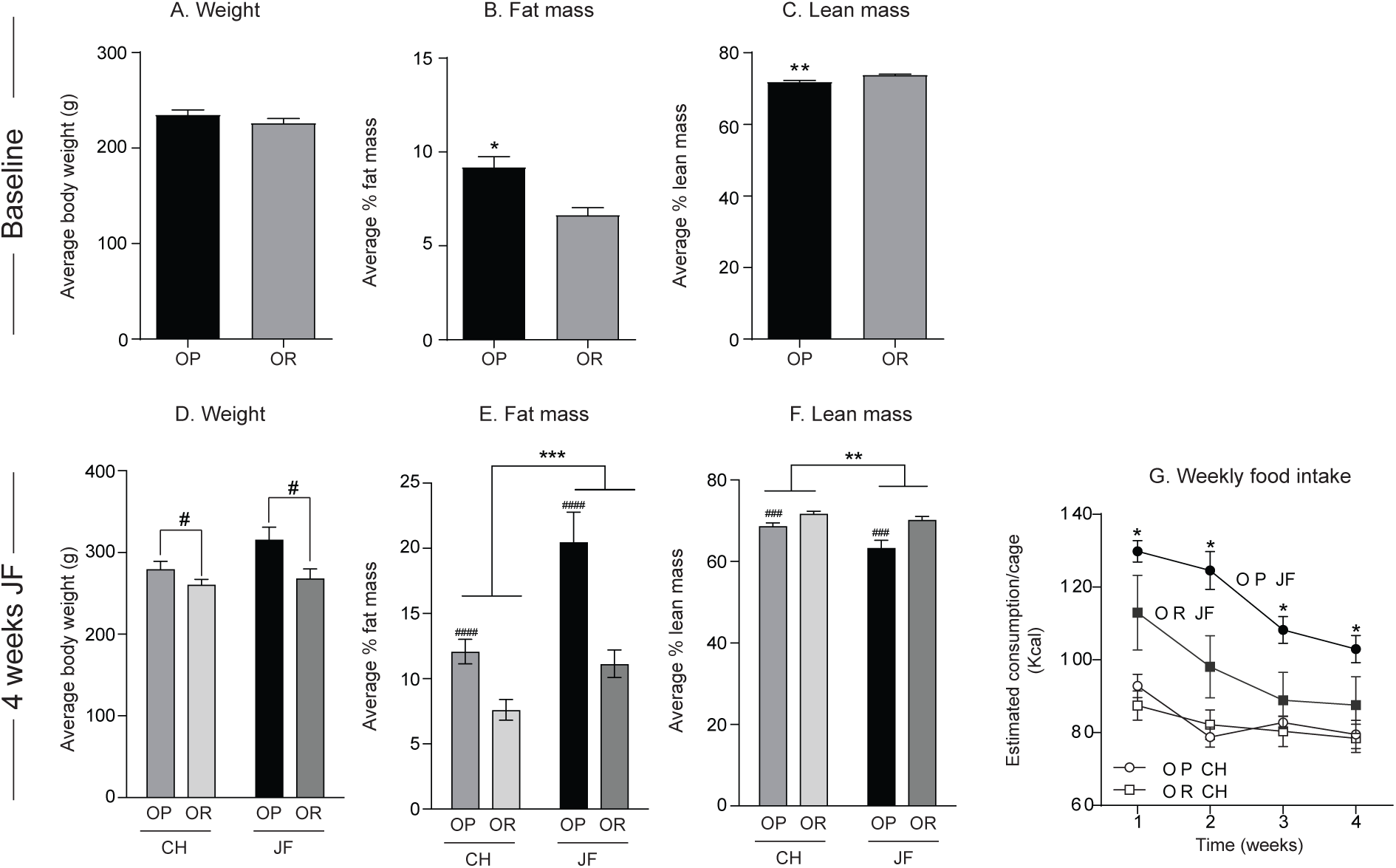
Verification of female obesity-prone and obesity-resistant phenotype. A-C) Weight and adiposity measures at baseline: Although body weight is similar between groups, obesity-prone rats (OP) have greater fat mass and lower lean mass vs. obesity-resistant (OR) rats. D-F) Weight and adiposity measures after 4 weeks of junk-food diet (JF) or chow (CH) consumption: Obesity-prone rats remain heavier than obesity-resistant rats, and have greater fat mass and lower lean mass vs. obesity-resistant rats. Junk-food increases fat mass and reduces lean mass in both groups, but the magnitude of this effect is stronger in obesity-prone rats. G) Home cage food intake during 4 weeks of junk-food or chow consumption: Junk-food consumption was greater than chow consumption in both groups, with obesity-prone rats consuming more junk-food than obesity-resistant rats. All data are shown as mean ±SEM unless otherwise noted. =differences between obesity-prone and obesity-resistant rats, *=differences between chow and junk-food, p<0.05.

Figure 1G shows food intake during the 4-week diet manipulation. As expected, junk-food consumption was greater than chow consumption in both groups, with obesity-prone rats consuming more junk-food than obesity-resistant rats (Fig. 1G: Three-way ANOVA main effect of time: *F*_(3,36)_=54.07, p<0.0001; main effect of diet: *F*_(1,12)_=22.5, p=0.0005; main effect of group: *F*_(1,12)_=4.33, p=0.06; and group x diet x time interaction: *F*_(3,36)_=3.31, p=0.03). In addition, chow consumption was similar between obesity-prone and obesity-resistant (Fig. 1G, open symbols) even though obesity-prone rats were heavier and had more fat mass than obesity-resistant rats. These data are consistent with previous reports in males (Vollbrecht et al., 2015) and confirm obesity-prone and obesity-resistant phenotypes in females.

### Experiment 1b: Food intake and motivation to obtain sucrose are reduced during estrus in obesity-prone and obesity-resistant females

Home cage food intake and responding during progressive ratio testing were examined in each phase of the cycle in naturally cycling females. Figure 2A shows example images capturing cell cytology distinctive of each phase of the cycle (20x magnification). Consistent with the existing literature, home cage food intake was reduced during the estrus phase compared to other phases of the cycle in both groups (Fig. 2B: Two-way RM ANOVA main effect of estrous phase *F*_(3,30)_=8.34, p<0.001; planned Sidak’s multiple comparisons proestrus vs. estrus, p<0.05). In addition, food intake was again greater in obesity-prone vs. obesity-resistant rats, regardless of estrous phase (Fig. 2B: Two-way RM ANOVA main effect of group: *F*_(1,10)_=28.40, p=0.0003). Importantly, there were no differences in number of cycles completed between obesity-prone vs. obesity-resistant groups across 13 days (data not shown).

**FIG. 2:**
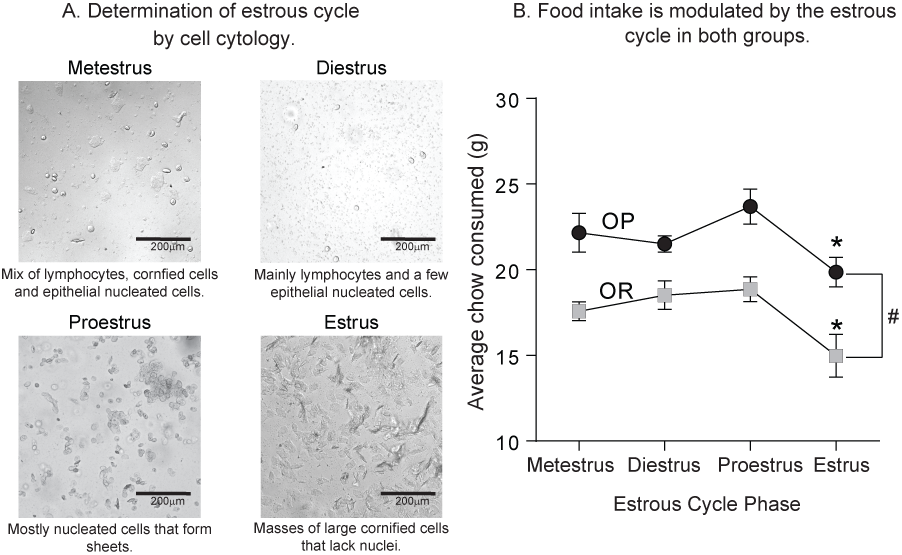
Home cage food intake decreases during estrus in both obesity-prone and obesity-resistant female rats. A) Representative pictures from vaginal lavages at each phase of the estrous cycle in female rats. B) Home cage chow consumption across the cycle. Homecage chow consumption was greater in obesity-prone (OP) vs. obesity-resistant (OR) rats, and was reduced during estrus in both groups. *=planned post-hoc comparisons, =obesity-prone vs. obesity-resistant rats, p<0.05.

We next determined how motivation to obtain sucrose changes across the cycle. Figure 3 shows behavior during initial fixed ratio training (Fig. 3A, B) and subsequent progressive ratio testing (Fig. 3C). Acquisition of instrumental responding for sucrose was similar between groups, with all rats preferentially responding on the active vs. inactive lever during FR1 (Fig. 3A: Three-way ANOVA main effect of lever *F*_(1,108)_=0.08, p<0.0001) and FR5 training (Fig. 3B: Three-way ANOVA main effect of lever *F*_(1,108)_=137.9, p<0.0001). In addition, the magnitude of responding was similar between obesity-prone and obesity-resistant groups. Progressive ratio testing was conducted across 8 consecutive days and analyzed by averaging break points within subjects (OP n = 10; OR n = 10) tested during metestrus/diestrus and proestrus/estrus. Break point was significantly lower during proestrus/estrus vs. metestrus/diestrus in both obesity-prone and obesity-resistant groups (Fig. 3C: Two-way RM ANOVA main effect of estrous cycle phase: *F*_(1,18)_=12.05, p<0.01). Although some visual trends were present, no significant differences in average break point between obesity-prone and obesity-resistant groups were observed (Fig. 3C: Two-way RM ANOVA effect of group: *F*_(1,18)_=2.0, p=0.2). These data show that despite the differences in home cage food intake between obesity-prone and obesity-resistant rats, motivation to obtain sucrose pellets is similar between groups. Additionally, the estrous cycle affects motivation to obtain food and food intake similarly in both groups.

**FIG. 3:**
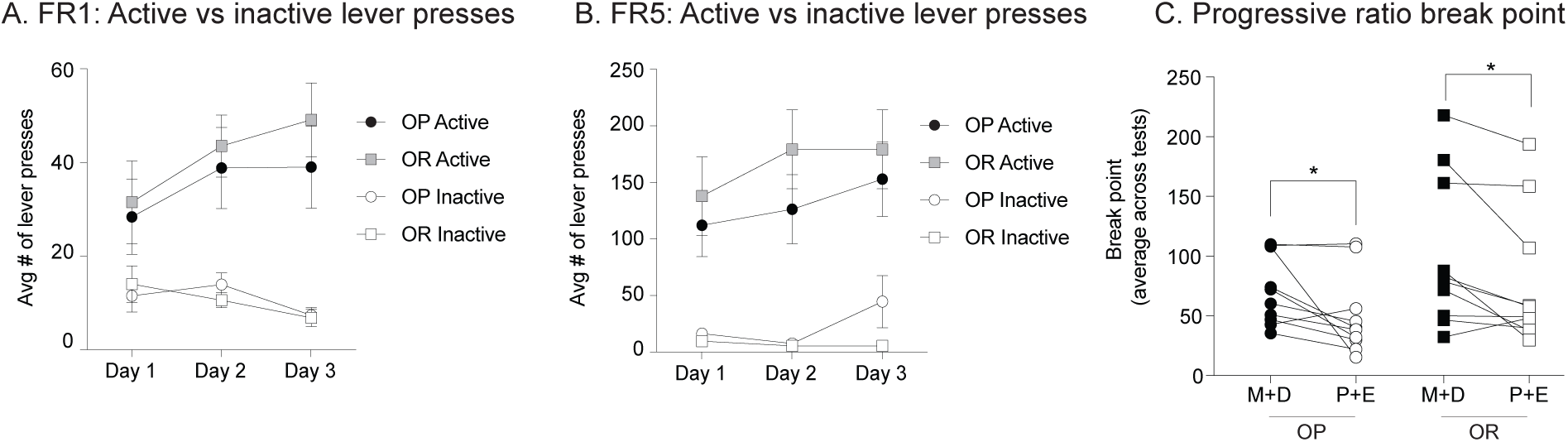
Motivation to work for a sucrose pellet decreases during proestrus/estrus in both obesity-prone and obesity-resistant rats. A, B) Average number of active and inactive lever presses across training. Acquisition of instrumental responding for sucrose was similar between groups. All rats preferentially responded on the active vs. inactive lever during fixed ratio 1 (FR1) and fixed ratio 5 (FR5) sessions. C) Average break point reached during progressive ratio testing. Break point was significantly lower during proestrus/estrus (P+E) vs. metestrus/diestrus (M+D) in both groups. No differences between obesity-prone (OP) and obesity-resistant (OR) groups were observed; *=p<0.05.

### Experiment 2: NAc core MSN intrinsic excitability is enhanced in obesity-prone vs. obesity-resistant rats during metestrus/diestrus

Recordings were made in both obesity-prone and obesity-resistant rats in metestrus/diestrus (Fig. 4 left graphs: OP n = 22 cells from 6 rats; OR n = 13 cells from 5 rats) or proestrus/estrus (Fig. 4 right graphs: OP n = 13 cells from 5 rats; OR n = 15 cells from 5 rats) to examine differences in MSN intrinsic excitability in the NAc core between groups and across the estrous cycle. Consistent with our previous results in males (Oginsky et al., 2016), the I/V curved showed a greater change in membrane potential in response to positive and negative current injections in MSNs from obesity-prone vs. obesity-resistant groups (Fig. 4C: Two-way RM ANOVA group x current injection interaction: *F*_(16,528)_=6.348, p<0.0001; main effect of group *F*_(1,33)_=5.683, p=0.02; main effect of current injection *F*_(16,528)_=263.9, p<0.0001; Sidak’s multiple comparison p<0.05). Similarly, during metestrus/diestrus, the number of action potentials elicited across current injections was greater in obesity-prone vs. obesity-resistant groups (Fig. 4D: Two-way RM ANOVA group x current injection interaction *F*_(14,462)_=3.04, p=0.0002; main effect of current injection *F*_(14,462)_=129.1, p<0.0001; main effect of group *F*_(1,33)_=3.67, p=0.06; Sidak’s multiple comparison p<0.05). In contrast, when comparisons were made during the proestrus/estrus phases, I/V relationships and the number of action potentials elicited by current injection were similar between groups (Fig. 4E, F).

**FIG. 4:**
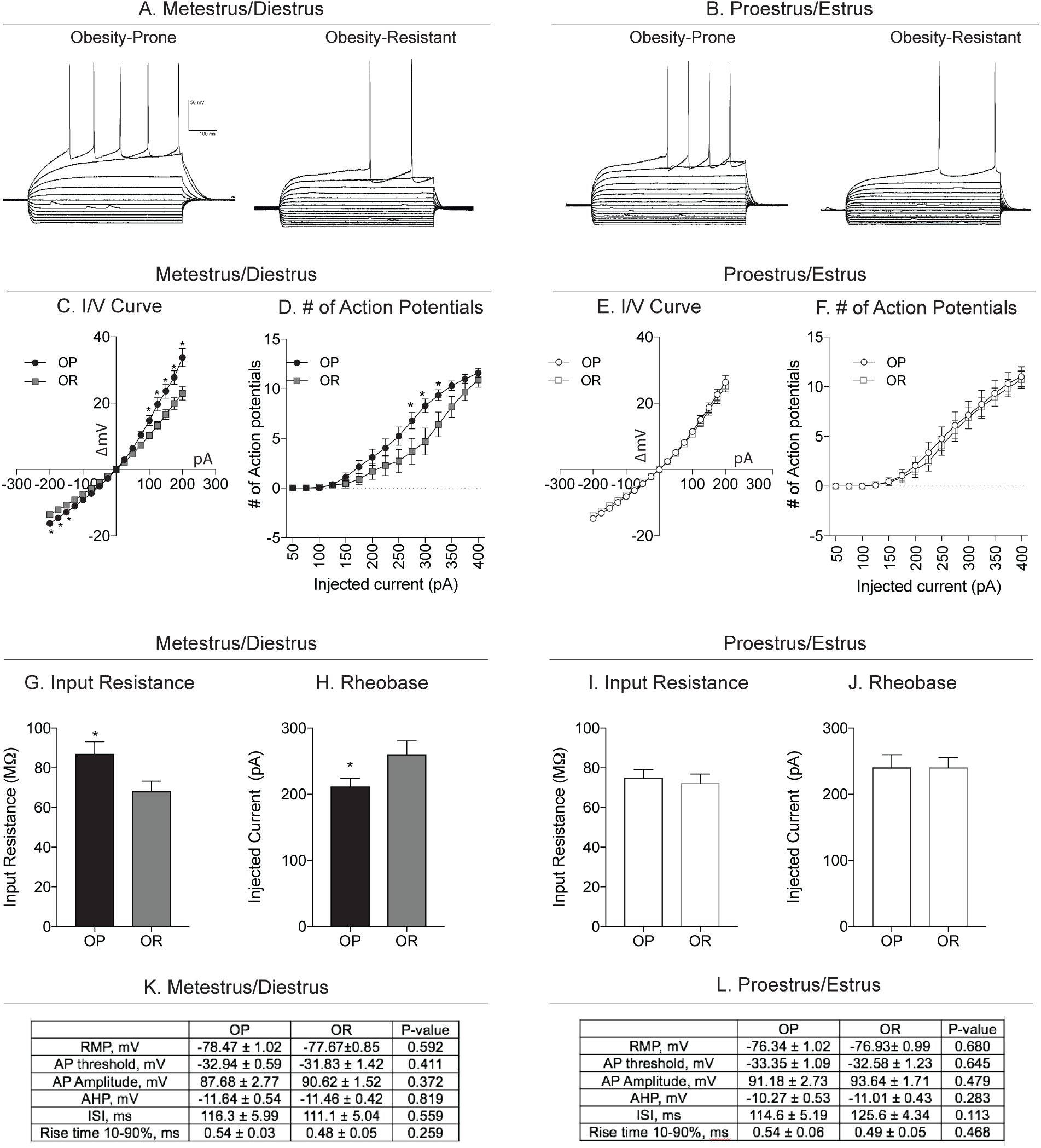
NAc core MSN intrinsic excitability is enhanced in obesity-prone vs. obesity-resistant rats during metestrus/diestrus. A) Example traces of current-clamp recordings from MSNs in slices from obesity-prone (OP) and obesity-resistant (OR) rats prepared during metestrus/diestrus. B) Example traces of current-clamp recordings from MSNs from obesity-prone and obesity-resistant rats prepared during proestrus/estrus. C, D) Current/voltage (I/V) relationship and number of action potentials in obesity-prone and obesity-resistant from slices made during metestrus/diestrus. I/V relationships are shifted to the right in in obesity-prone vs. obesity-resistant rats and the same current injection intensity elicits more action potentials in MSNs from obesity-prone vs. obesity-resistant rats. E, F) Current/voltage (I/V) relationship and number of action potentials in obesity-prone and obesity-resistant from slices made during proestrus/estrus. I/V relationships and the number of action potentials fired are similar between obesity-prone and obesity-resistant rats in proestrus/estrus. G, H) Input resistance and rheobase: Obesity-prone rats have a higher input resistance and lower rheobase vs. obesity-resistant rats. I, J) Input resistance and rheobase during proestrus/estrus: No differences were observed in input resistance or rheobase between obesity-prone and obesity-resistant rats. K, L) Cell parameters from recordings conducted in slices prepared during metestrus/diestrus and proestrus/estrus. No differences in resting membrane potential (RMP), action potential threshold, action potential amplitude (AP), afterhyperpolarization amplitude (AHP), inter-spike interval (ISI) or rise time were observed between groups or cycle phase metestrus/diestrus or proestrus/estrus; *=p<0.05.

Consistent with group differences found during metestrus/diestrus, input resistance was greater in obesity-prone rats (Fig. 4G: Two-tailed unpaired t-test, *t*_(33)_=2.09, p=0.04) and rheobase was reduced (Fig. 4H: Two-tailed unpaired t-test, *t*_(33)_=2.15, p=0.04) compared to obesity-resistant rats during metestrus/diestrus. However, these group differences were absent when comparison were made during the proestrus/estrus phases (Fig. 4I,J). We did not observe differences in basal cell parameters including resting membrane potential (RMP), action potential threshold, amplitude, rise time (10-90%), after-hyperpolarization (AHP) amplitude, or duration of the first interspike interval between recordings made during metestrus/diestrus in obesity-prone and obesity-resistant rats (Fig. 4K) or proestrus/estrus (Fig. 4L) phases. In sum, these data show that similar to male obesity-prone rats, basal intrinsic excitability is enhanced in female obesity-prone vs. obesity-resistant rats, but these differences are only apparent when recordings are made during the metestrus/diestrus phase of the estrous cycle.

### Experiment 3: Cue-triggered motivation is modulated by the estrous cycle in obesity-prone, but not obesity-resistant rats

Here, we determined how cue-triggered motivation in the form of conditioned approach varies across the cycle in naturally cycling obesity-prone and obesity-resistant rats. Figure 5 shows behavior during initial conditioning (Fig. 5A-C) and subsequent testing in extinction conditions (Fig. 5D). Behavior during initial Pavlovian conditioning was similar between groups, with both obesity-prone and obesity-resistant rats acquiring a similar magnitude of discrimination between the CS+ and CS-(Fig. 5A OP: Two-way RM ANOVA: main effect of CS *F*_(1,30)_=9.79 p<0.01; Fig. 5B OR: Two-way RM ANOVA: main effect of CS *F*_(1,30)_=10 p<0.01; Fig. 5C OP vs. OR: main effect of session: *F*_(4,240)_=4.98, p<0.001; no effect of group *F*_(1,60)_=0.17, p=0.7; no group x session interaction *F*_(4,120)_=1.78, p=0.14). Rats were then tested for conditioned approach in extinction conditions and data were analyzed according to which phase of the cycle rats were in during testing (Fig. 5D). The CS+ elicited strong conditioned approach in all groups (Fig. 5D: Two-way RM ANOVA period (ITI, CS+,CS-) x group interaction *F*_(6,116)_=3.1, p=0.007; main effect of group *F*_(3,58)_=3.85, p=0.01; main effect of period *F*_(2,116)_=222.7, p<0.0001; Sidak’s multiple comparison: CS+ vs. ITI and CS+ vs. CS-, P<0.0001). However, the magnitude of conditioned approach in response to the CS+ was significantly greater in obesity-prone rats tested during metestrus/diestrus compared to all other groups (Fig. 5D; Sidak’s multiple comparison, p<0.05). Furthermore, differences in CS+ responding between obesity-prone and obesity-resistant groups were driven entirely by enhanced approach behavior during metestrus/diestrus vs. proestrus/estrus in obesity-prone, but not obesity-resistant rats (Fig. 5D: Sidak’s multiple comparison CS+: OP-M+D vs. OP-P+E, p=0.0001; CS+: OR-M+D vs. OR-P+E, p=0.73). No significant differences in responding during the ITI or CS-were found between groups or across cycle phase. Thus, in obesity-prone, but not obesity resistant rats, cue-triggered motivation varied across the cycle and was stronger in obesity-prone vs. obesity-resistant rats tested in metestrus/diestrus phases. Taken with results from progressive ratio testing above, data show a dissociation between the ability of the cycle to influence motivation for food (Fig. 3) but not motivation triggered by a food cue in obesity-resistant rats (Fig. 5D).

**FIG. 5:**
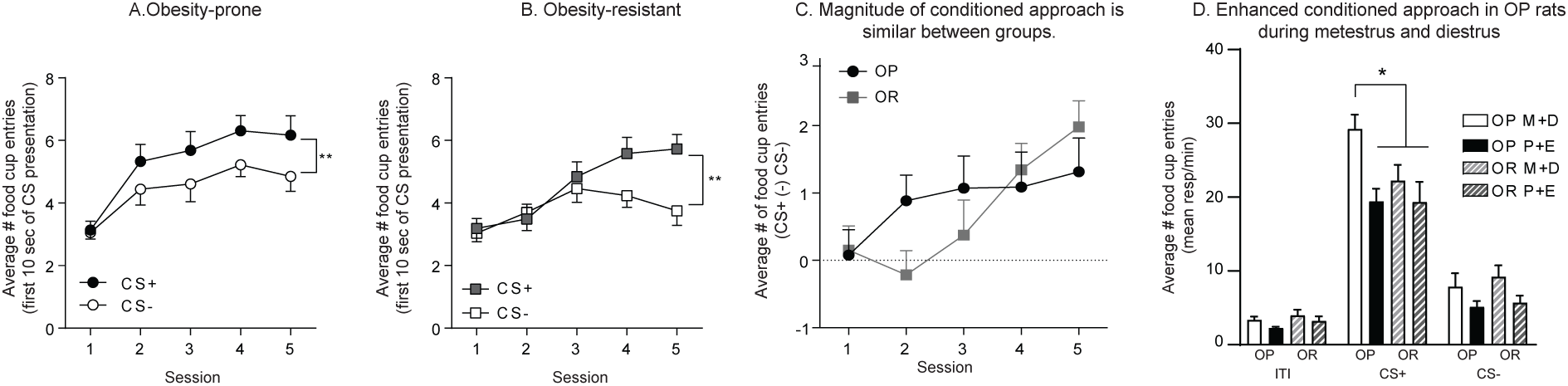
Cue-triggered motivation is modulated by the estrous cycle in obesity-prone but not in obesity-resistant rats. A) Average number of food cup entries during the first 10 seconds of CS presentation in obesity-prone rats (OP). Obesity-prone rats discriminate between the CS+ and the CS-during initial Pavlovian conditioning. B) Average number of food cup entries during the first 10 seconds of CS presentation in obesity-resistant rats (OR). Obesity-resistant rats discriminate between the CS+ and the CS-during initial Pavlovian conditioning. C) Average difference in the number of food cup entries during the first 10 sec of CS+ vs. CS-presentation. No difference in magnitude of conditioned approach between obesity-prone and obesity-resistant rats were observed. D) Extinction Test: Average number of food cup entries during the CS+, CS- and intertrial interval (ITI) in obesity-prone and obesity-resistant rats tested in metestrus/diestrus (M+D) and proestrus/estrus (P+E). Conditioned approach is stronger during metestrus/diestrus vs proestrus/estrus in obesity-prone, but not obesity-resistant rats. In addition, the magnitude of conditioned approach is greater in obesity-prone vs. obesity-resistant rats tested during the metestrus/diestrus phase. *=p<0.05, **=p<0.01.

### Experiment 4: Repeated treatment with estradiol and progesterone is sufficient to decrease cue-triggered motivation in obesity-prone rats

Given that the estrous cycle modulates cue-triggered motivation in obesity-prone but not obesity-resistant rats (Exp. 3), we next determined whether single vs. repeated treatment with estradiol and progesterone in ovariectomized rats is sufficient to replicate this effect in obesity-prone rats (Fig. 6A: experimental timeline). As expected, obesity-prone rats learned to discriminate between the CS+ and the CS-during Pavlovian training (Fig. 6B: Two-way RM ANOVA main effect of CS *F*_(1,19)_=25.6, p<0.0001; significant session × CS interaction *F*_(7,133)_=3.74, p=0.001; Sidak’s post-test p<0.05). A single cycle of estradiol and progesterone treatment was not sufficient to alter conditioned approach behavior compared to vehicle treated controls (Fig. 6C). However, repeated estradiol and progesterone treatment (repeated E+P), decreased conditioned approach compared to controls (Fig. 6D: Two-way RM ANOVA; main effect of treatment, p<0.05; significant period (ITI, CS+ and CS-) × treatment interaction, p <0.05; Sidak’s multiple comparisons CS+: vehicle vs. hormone treatment, p=0.002). It should be noted that the same rats were tested twice (see timeline Fig. 6A). Thus, it is not surprising that the magnitude of conditioned approach was lower overall during the second test conducted under extinction conditions (Fig. 6D vs. C). Nonetheless, these data show that single hormone treatment does not affect conditioned approach, whereas repeated hormone treatment reduces it in obesity-prone rats.

**FIG. 6:**
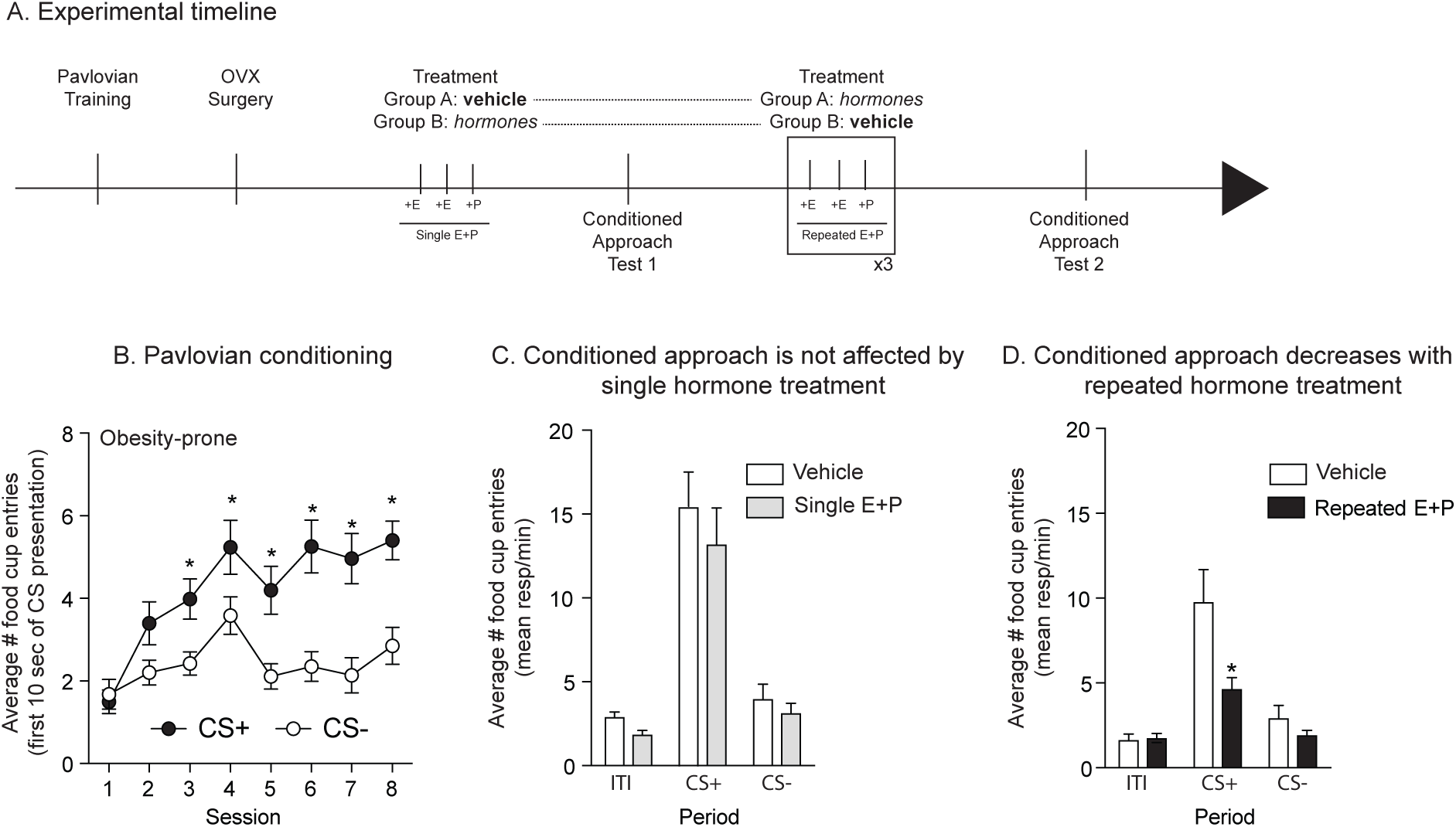
Repeated treatment with estradiol and progesterone decreased conditioned approach in obesity-prone females. A) Schematic of experimental timeline. Rats received 8 sessions of Pavlovian conditioning, followed by ovariectomy surgery (OVX) and 10 days of recovery before being treated with estradiol (+E) and progesterone (+P) or vehicle. 4-6 hours after the last progesterone injection rats were tested in extinction conditions. B) Average number of food cup entries during the first 10 seconds of CS presentation in obesity-prone rats (OP). Obesity-prone rats learned to discriminate between the CS+ and CS-. C) Average number of food cup entries during the CS+, CS- and inter-trial interval (ITI) in vehicle and hormone treated groups. A single cycle of hormone treatment is not sufficient to decrease conditioned approach. D) Average number of food cup entries during the CS+, CS- and intertrial interval (ITI) in vehicle and hormone treated groups. Repeated estradiol and progesterone treatment decreased conditioned approach compared to vehicle treated controls, although repeated testing reduced the magnitude of conditioned approach in both treatment groups. *= p<0.05.

### Experiment 5a: Repeated treatment with estradiol and progesterone is sufficient to decrease cue-triggered motivation in outbred rats

Here, we determined if the decrease in cue-triggered motivation with repeated E+P observed in ovariectomized obesity-prone rats also extend to ovariectomized outbred Sprague-Dawley females (see Fig. 7A for experiment timeline). As expected, outbred females learned to discriminate between the CS+ and the CS-during Pavlovian conditioning (Fig. 7B: Two-way RM ANOVA: main effect of CS *F*_(1,19)_=13.83, p=0.002; significant session × CS interaction *F*_(7,133)_=7.78, p<0.0001, Sidak’s multiple comparisons p<0.05). Consistent with effects in selectively-bred rats, repeated estradiol and progesterone treatment resulted in a decrease in conditioned approach behavior to the CS+ (Fig. 7C: Two-way RM ANOVA main effect of treatment p<0.05; significant period × treatment interaction p=0.05; Sidak’s multiple comparisons CS+: vehicle vs. hormone treated, p<0.05). There was a slight effect of hormone treatment on food cup entries during the ITI (Fig. 7C; vehicle vs. hormone treated p=0.056) but not during the CS-(Fig. 7C vehicle vs. hormone treated p=0.83). However, when comparing across all three separate experiments (see below), there do not appear to be consistent effects of hormone replacement on responding during the ITI. In sum, data from outbred females replicate the primary effect of hormone treatment found in obesity-prone rats, supporting a role for ovarian hormones in reductions in conditioned approach behavior during metestrus/diestrus.

**FIG. 7:**
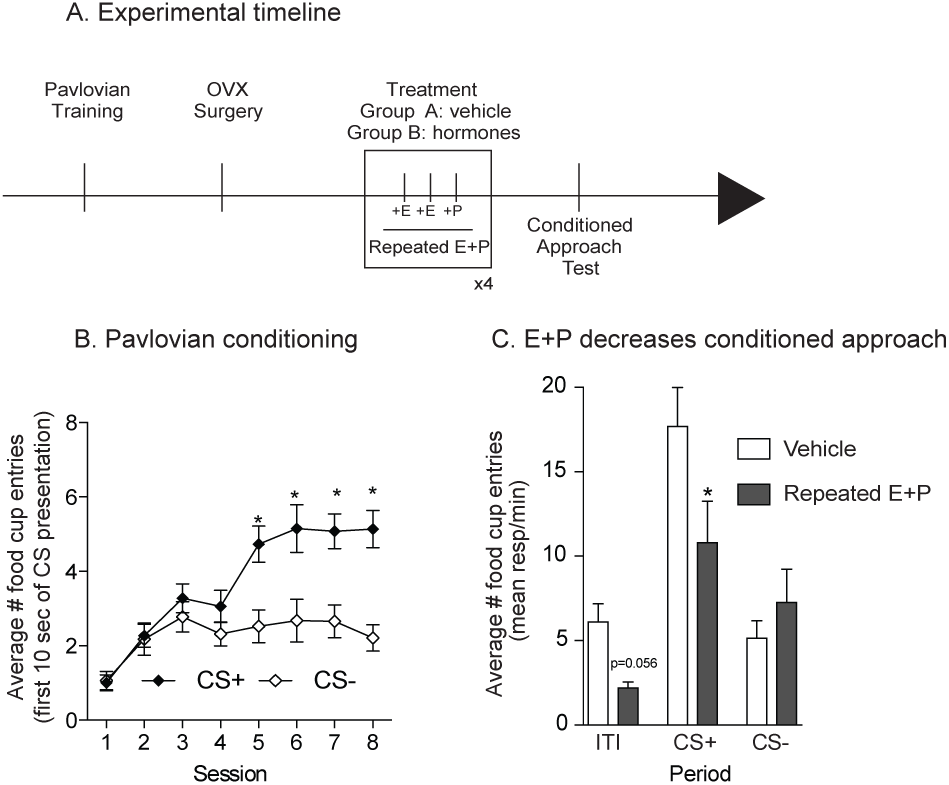
Repeated treatment with estradiol and progesterone decreased conditioned approach in outbred females. A) Schematic of experimental timeline. Rats received 8 sessions of Pavlovian conditioning, followed by ovariectomy surgery (OVX) and 10 days of recovery before being treated with estradiol (E) and progesterone (P) or vehicle. 4-6 hours after the last progesterone injection rats were tested in extinction conditions. B) Average number of food cup entries during the first 10 seconds of CS presentation. Outbred rats learned to discriminate between the CS+ and the CS-. C) Average number of food cup entries during the CS+, CS- and intertrial interval (ITI) in vehicle and hormone treated groups. Four cycles of estradiol and progesterone treatment decreased conditioned approach in outbred rats. *=p <0.05.

### Experiment 5b: Estradiol and progesterone act synergistically to decrease cue-triggered motivation in outbred rats

Above we demonstrate that repeated estradiol and progesterone treatment decreases cue-triggered motivation in both obesity-prone and outbred females. Here we evaluated the ability of repeated estradiol or progesterone alone to reduce cue-triggered motivation in outbred rats (see Fig. 8A for experimental timeline). We also included an additional experimental group that was given repeated treatment with both hormones in order to replicate effects above in a separate group of rats. After acquisition of Pavlovian discrimination (Fig. 8B: main effect of CS *F*_(1,38)_=26.9, p<0.0001; significant session × CS interaction *F*_(7,266)_=6.29, p<0.0001, Sidak’s multiple comparisons, p<0.05), rats were separated into 4 treatment groups [vehicle (veh), estradiol and progesterone (E+P), estradiol alone (E) and progesterone alone (P)], counter-balanced for behavior during the last Pavlovian training session.

**FIG. 8:**
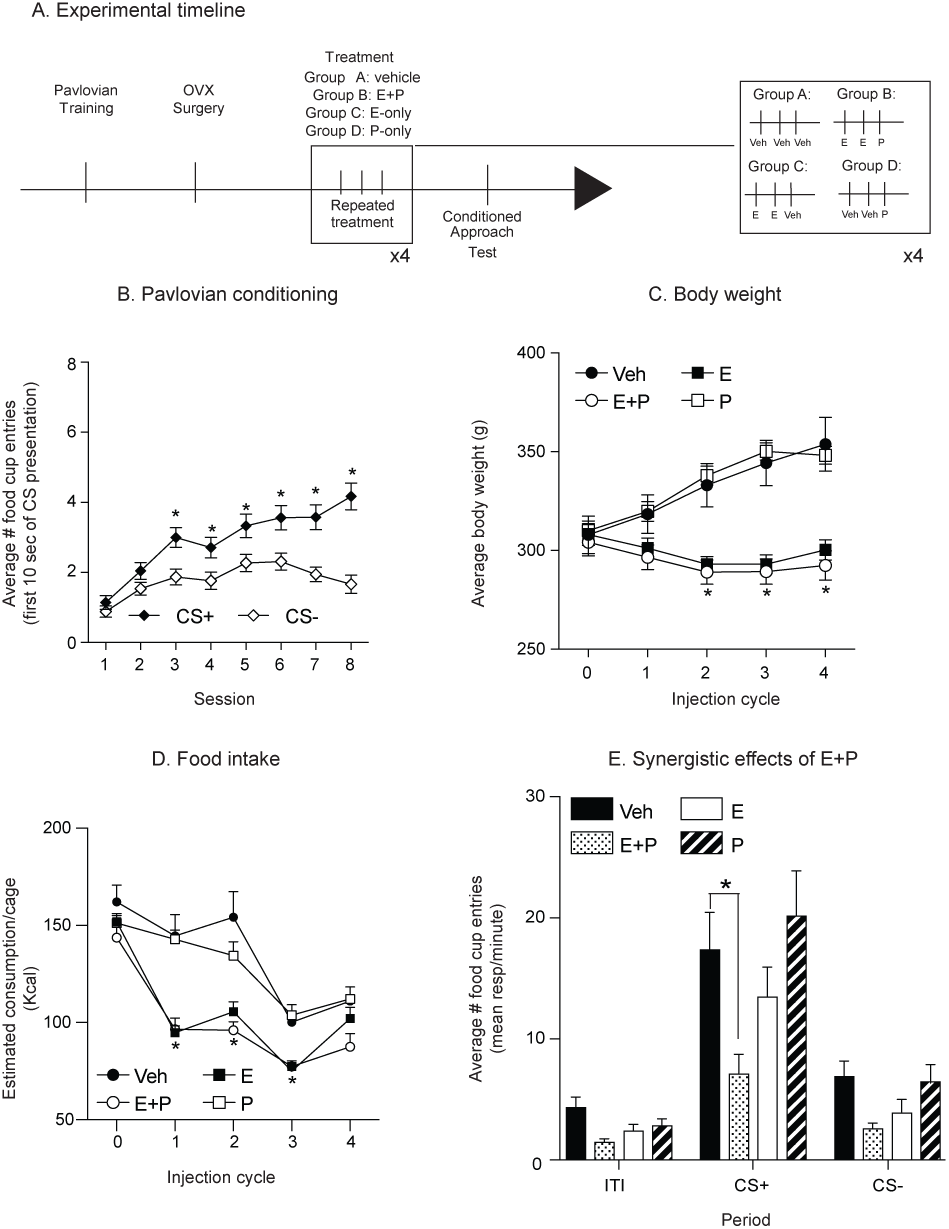
Estradiol and progesterone act synergistically to decrease cue-triggered motivation in outbred rats. A) Schematic of experimental timeline. Outbred rats received 8 sessions of Pavlovian conditioning, followed by ovariectomy surgery (OVX) and 10 days of recovery. Rats were the counterbalanced by weight and divided into 4 groups: vehicle, estradiol and progesterone (E+P), estradiol alone (E) and progesterone alone (P). B) Average number of food cup entries during the first 10 seconds of CS presentation. Outbred rats learned to discriminate between the CS+ and the CS-. C, D) Average body weight and food intake across the treatment period. Estradiol treatment (E or E+P) reduced body weight and food intake compared to the vehicle (Veh) and progesterone (P) groups. E) Average number of food cup entries during the CS+, CS- and intertrial interval (ITI) in vehicle and hormone treated groups. Estradiol or progesterone alone did not affect conditioned approach. However, repeated estradiol and progesterone treatment decreased conditioned approach in outbred rats. Thus, although estradiol is sufficient to reduce weight and food intake, both estradiol and progesterone are needed to reduce conditioned approach. *=p<0.05.

Throughout hormone treatment we measured body weight and food intake. We found that body weight and food intake were reduced in both estradiol treated groups (E and E+P; Fig. 8C body weight: Two-way RM ANOVA: main effect of injection cycle *F*_(4,144)_=45.63, p<0.0001; main effect of group *F*_(3,36)_=8.43, p=0.0002; significant interaction injection cycle x group *F*_(12,144)_=45.21, Sidak’s multiple comparisons p<0.05; Fig. 8D food intake: Two-way RM ANOVA: main effect of injection cycle *F*_(4,64)_=70.14, p<0.0001; main effect of group *F*_(3,16)_=17.88, p <0.0001; significant injection cycle x group *F*_(12,64)_=3.73, p=0.0003, Sidak’s multiple comparison, p <0.05). Consistent with experiment 5a above, repeated E+P treatment significantly decreased conditioned approach to the CS+ (Fig. 8E: Two-way RM ANOVA significant period x treatment interaction *F*_(6,68)_=2.6; main effect of period *F*_(2,68)_=62.5, p<0.0001; main effect of treatment *F*_(3,34)_=5.4, p=0.004; Planned Sidak’s multiple comparisons CS+: vehicle vs. E+P, p=0.0008), with no effect of treatment on responding during the CS- or ITI (vehicle vs. E+P; p=0.84, 0.46 respectively). However, treatment with estradiol or progesterone alone were not sufficient to decrease conditioned approach in outbred rats (Fig. 8E: Sidak’s multiple comparisons CS+: Veh vs. E, p=0.54; Fig. 8E: Sidak’s multiple comparisons CS+: Veh vs. P, p=0.84). Together these data suggest a synergistic effect of repeated estradiol and progesterone treatment on cue-triggered motivation, even though estradiol is sufficient to decrease home-cage food intake and body weight.

## IV. DISCUSSION

Naturally occurring alterations in estradiol influence food intake in females. However, how motivational responses to food cues are affected by the estrous cycle or elevations in ovarian hormones is unknown. In addition, while individual susceptibility to obesity is accompanied by enhanced incentive motivational responses to food cues and increased intrinsic excitability of MSNs within the NAc of males, studies in females are lacking (see Introduction and Alonso-Caraballo et al., 2018). Here, we show that intrinsic excitability of NAc MSNs and conditioned approach behavior are enhanced in female obesity-prone vs. obesity-resistant rats when measured during metestrus/diestrus. However, neural and behavioral responses between these groups were similar when measured during proestrus/estrus. The emergence of group differences during metestrus/diestrus were due to effects of the cycle on NAc excitability and cue-triggered motivation within obesity-prone, but not obesity-resistant rats. Additionally, we found that estradiol and progesterone treatment in ovariectomized females reduced conditioned approach behavior in obesity-prone and outbred Sprague-Dawley rats. To our knowledge, these data are the first to demonstrate cycle- and hormone-dependent effects on the motivational response to a food cue, and the only studies to date to determine how individual susceptibility to obesity influences NAc excitability, cue-triggered food-seeking, and differences in the regulation of these neurobehavioral responses by the cycle.

### Effects of the cycle on weight, food intake, and motivation for sucrose in obesity-prone and obesity-resistant females

We began by verifying obesity-prone and obesity-resistant phenotypes in females. As expected, female obesity-prone rats were heavier, ate more, and had greater fat mass and less lean mass than obesity-resistant females (Fig. 1). In addition, weight gain induced by a junk-food diet was more pronounced in obesity-prone vs. obesity-resistant females. Instrumental responding for a sucrose pellet during FR1, FR5 or progressive ratio testing was similar between groups (Fig. 3), despite differences in home cage food consumption. In males break point is modestly elevated in obesity-prone vs. obesity-resistant rats working for sucrose (Vollbrecht et. al., 2015). In the current study, pressing on the active lever resulted in the delivery of a sucrose pellet but no discrete cue. However, in our previous study of males, active lever presses resulted in the presentation of an auditory cue in addition to the delivery of the sucrose pellet. Thus, its possible that inclusion of a discrete cue added salience to the motivation to work for food in our previous study using males, and that the absence of cue presentation in the current study contributed to similar break points in female obesity-prone and resistant groups. However, the absence of strong differences in break point between obesity-prone and obesity-resistant females here did not impede our ability to observe effects of the cycle on this behavior, discussed further below. Overall, obesity-prone and -resistant phenotypes of females are similar to those previously reported for males of these selectively bred lines (Vollbrecht et al., 2015, 2016). Additionally, within naturally cycling obesity-prone and obesity-resistant females, food intake and body weight were reduced during estrus (Fig. 2B). Furthermore, motivation to obtain sucrose was lower when rats were tested during proestrus/estrus vs. metestrus/diestrus (Fig. 3C), regardless of susceptibility to obesity. These changes in motivation to obtain sucrose in naturally cycling females are consistent with variations in home cage food intake across the cycle found here and with established roles for ovarian hormones in the regulation of food intake (Tarttelin Gorski, 1971; Asarian and Geary, 2013).

### Differences in NAc MSN intrinsic excitability in obesity-prone vs. obesity-resistant rats

The NAc is comprised predominantly of MSNs which integrate both dopaminergic and glutamatergic inputs to ultimately influence behavioral responses to reinforces like food, sex, and drugs as well as cues paired with them (Schultz, 1997; Berridge and Robinson, 1998; Schultz, 2013; Wolf et al., 2002; Wolf, 2003). One factor that strongly influences the output of the NAc is the intrinsic excitability of MSNs, i.e., how readily they can be depolarized and fire action potentials (Nicola et al., 2000; Hu, 2007). We found that excitability of MSNs in the NAc core is enhanced in obesity-prone vs. obesity-resistant females when recordings are made during metestrus/diestrus, but that this group difference is not apparent when recordings are made during proestrus/ estrus (Fig. 4). This was due to cycle effects on MSN excitability in obesity-prone, but not obesity-resistant rats. Specifically, in obesity-prone rats changes in voltage at positive current injections were significantly greater during metestrus/diestrus vs. proestrus/estrus. This is similar to effects of the cycle on NAc excitability in outbred Sprague-Dawley females (Proao et al., 2018). To our knowledge this is the only other study to examine effects of the estrous cycle on MSN excitability, and specific mechanisms by which ovarian hormones influence NAc excitability have not been determined, although there is evidence for sex differences (Cao et al., 2018).

MSN excitability is largely determined by the number and distribution of voltagegated potassium channels, with inwardly-rectifying *K*^+^ currents (I_*KIR*_) dominating at negative membrane potentials (< −90mV) and A-type *K*^+^ currents (I_*A*_) dominating at positive membrane potentials (*>* +40mV; Nisenbaum and Wilson 1995; Perez et al. 2006). Differences during metestrus/diestrus between obesity-prone and obesity-resistant females were present across a wide range of current injections, but were most pronounced at positive potentials. This, in combination with a lower rheobase and absence of differences in resting membrane potential, suggests that group differences may be due to lower IA in obesity-prone vs. obesity-resistant females. These basal group differences in excitability during metestrus/diestrus are similar to those seen in male obesity-prone vs. obesity-resistant rats where differences were found across the I/V curve and were accompanied by a lower rheobase and an absence of differences in resting membrane potential (Oginsky et al., 2016).

The reduction in excitability during proestrus/estrus vs. metestrus/diestrus within obesity-prone rats is consistent with the ability of estradiol to enhance *I*_*A*_, thereby reducing excitability in cultured hippocampal neurons (Zhang et al., 2015). Progesterone also increases during the proestrus phase, and progesterone can block voltage-gated sodium and calcium channels, resulting in decreased neuronal excitability (Kelley Mermelstein, 2011). However, we did not observed changes in the threshold for action potential firing or action potential rise time, which are mediated by *Na*^2+^ currents, nor did we find changes in after hyperpolarization amplitude, which are mainly influenced by *Ca*^2+^ channels. Thus, it seems less likely that alterations in excitability observed across the cycle in obesity-prone rats are mediated by progesterone. Of course, many hormones change across the cycle, and effects observed here could be either direct, or indirect. Nonetheless, these data demonstrate that in the NAc core MSN excitability is enhanced in obesity-prone rats during phases of the cycle when motivational responses to food cues are also elevated (see below for additional discussion of this relationship).

### Basal differences in cue-triggered motivation in obesity-prone vs. obesity-resistant females

Food intake is not only regulated by homeostatic feedback, but external signals including food cues also influence the motivation to eat independent of hunger state (Derman Ferrario, 2018; Holland, 1977). These cue-triggered urges to seek out and consume food contribute to opportunistic eating that drives obesity (Ferrario et al., 2016, Ferrario, 2017, Stice et al., 2013) and are more pronounced in male obesity-susceptible vs. -resistant populations (Robinson et al., 2015, Derman Ferrario, 2018, Alonso-Caraballo et al., 2018). Conditioned approach behavior was stronger in female obesity-prone vs. obesity-resistant rats (Fig. 5D). This is consistent with data from males, where the magnitude of Pavlovian conditioned approach, conditioned reinforcement, and Pavlovian to instrumental transfer are greater in obesity-susceptible vs. resistant populations (Robinson et al., 2015; Derman Ferrario, 2018). Taken as a whole, these data support the idea enhanced responsivity to the motivational properties of food cues is likely a phenotypic, neurobehavioral difference in those individuals that are more vs. less susceptible to diet-induced weight gain.

Difference in conditioned approach between obesity-prone and obesity-resistant females was apparent when naturally cycling rats were tested during the metestrus/diestrus phases of the cycle, but the magnitude of conditioned approach was similar across groups when tested during proestrus/estrus. Specifically, in obesity-prone rats conditioned approach was elevated during metestrus/diestrus compared to proestrus/estrus (Fig. 5), consistent with shifts in home cage food intake and break point across the cycle. In contrast, in obesity-resistant females, conditioned approach behavior was stable across the cycle, even though motivation for food itself and food consumption rose and fell with the cycle in a manner comparable to that found in obesity-prone rats (Fig. 5). This dissociation between motivation for food and motivational responses triggered by a food cue does not appear to be due to deficits in Pavlovian learning in obesity-resistant rats, as behavior during acquisition and the degree of discrimination between the CS+ and CS-were similar to that seen in the obesity-prone group (Fig. 5). Indeed, obesity-resistant rats show quite reliable conditioned approach and discrimination between the CS+ and CS-during testing. Thus, the lack of cycle dependent shifts in conditioned approach, despite shifts in food intake and break point, suggest a potential mechanistic dissociation between appetitive and consummatory behaviors (see Berridge et al., 2010 for review). Although speculative, given the role of the NAc core in motivational responses to food cues (Kelley et al., 2005; Aitken et al., 2016) its possible that the cycle-dependent fluctuations in MSN excitability in obesity-prone rats discussed above, could contribute to differences in the modulation of conditioned approach across the cycle in obesity-prone but not obesity-resistant rats. This should be examined in future studies.

### Ovarian hormones modulate cue-triggered motivation in obesity-prone and outbred females

Given that the magnitude of conditioned approach varied across the cycle in intact obesity-prone rats, we also determined the degree to which estradiol and progesterone treatment of ovariectomized rats reduces conditioned approach behavior. A single cycle of estradiol and progesterone treatment did not alter the expression of conditioned approach behavior (Fig. 6C), whereas repeated cycles of this treatment were sufficient to reduce conditioned approach in three separate experiments (Figs. 6D, 7C, 8E). Importantly, this effect was found in both obesity-prone and outbred Sprague Dawley females; thus, it is not unique to our selectively bred rat strain. Similarities across obesity-prone and outbred rats are not surprising given that obesity-susceptible rats can be identified in the outbred population (e.g., Levin et al., 1997; Robinson et al., 2015; Madsen et al., 2010).

The fact that repeated hormone treatment was needed in order to recapitulate effects seen in naturally cycling rats may in part be due to effects of ovariectomy. Following Pavlovian conditioning rats were ovariectomized and hormone replacement did not begin until rats failed to enter estrus within an 8-10 day period. The removal of the ovaries could itself produce neuroadaptations, such as changes in progesterone and estradiol receptor expression. Thus, repeated cycles of estradiol and progesterone may have been needed in order to compensate for adaptations induced by ovariectomy itself. Nonetheless, these data demonstrate that ovarian hormones strongly influence conditioned approach elicited by a food cue. Furthermore, repeated treatment with estradiol or progesterone alone were not sufficient to reduce conditioned approach behavior (Fig. 5). In contrast, repeated treatment with estradiol was sufficient to decrease home cage food intake and body weight, consistent with previous reports (Blaustein Wade, 1975). However, it was only when estradiol and progesterone were co-administered that a decrease in conditioned approach was observed. This again points to dissociable mechanisms through which ovarian hormones regulate food intake vs. food-seeking.

To our knowledge, this is the first demonstration that ovarian hormones influence cue-triggered food-seeking. There have been many studies examining how ovarian hormones influence motivation for primary reinforcers like food, sex, and potentially addictive substances like cocaine (Yoest et al., 2018; Yoest et al., 2014; Rivera Stincic, 2018; Frye, 2007). Although the mechanisms underlying these behaviors are not well understood, ovarian hormones do influence both dopaminergic and glutamatergic transmission within the NAc core in ways that are consistent with the behavioral effects found here (Micevych Meisel, 2017, Tonn Eisinger et al., 2018, Yoest et al., 2018). In addition to effects of dopamine on NAc intrinsic excitability (discussed above), estradiol treatment also decreases dendritic spine density (an indirect measure of excitatory synapses) in the NAc (Peterson et al., 2015), and reduces AMPA receptor specific binding (Cyr et al., 2001) and GluA2 AMPAR subunit mRNA in the NAc (Le Saux et al., 2006). Thus, reductions in the motivational responses to food cues may be related to the effects of estradiol on excitatory transmission in the NAc. However, it’s important to keep in mind that effects on conditioned approach required both estradiol and progesterone treatment, suggesting a more complex mechanism.

### Conclusions and Future Directions

In sum, we find that in females individual susceptibly to obesity is associated with enhanced motivational responses to food cues and increased intrinsic excitability of NAc core MSNs, consistent with overall patterns found in males. In addition, both estradiol and progesterone are needed to reduce conditioned approach behavior in obesity-prone rats, whereas estradiol treatment alone is sufficient to reduce food intake and motivation for food. These data suggest dissociations between hormonal regulation of food-seeking vs. consumption. Furthermore, interactions between individual susceptibility to obesity and the regulation of incentive motivation by ovarian hormones likely represent phenotypic differences that may be mediated by altered NAc responsivity. Future studies of the precise mechanisms involved, as well as the effects of diet-induced obesity and consumption of sugary, fatty foods are needed in order to understand the fundamental neurobiological mechanisms of food-seeking in the non-obese and obese state. Finally, effects of ovarian hormones on conditioned approach were similar in obesity-prone and outbred females, suggesting that neurobehavioral effects of ovarian hormones observed here extend to outbred populations.

## Acknowledgments

This work was supported by R01DK106188-02-S1 and 1F99NS108549-01 to YAC, and R01DK106188; 1R01DK115526-01, and R21DA045277 to CRF. Additionally, we thank Ms. Sophia Dunlap, Ms. Limmy Kim and Ms. Liana Dunietz for their help with behavioral portions of this study and Dr. Johnny Sexton for the use of his light microscope.

## Author Contributions

YAC designed, conducted experiments, analyzed data, and wrote the manuscript. CRF designed experiments, analyzed data, and wrote the manuscript.

